# Generative whole-brain dynamics models from healthy subjects predict functional alterations in stroke at the level of individual patients

**DOI:** 10.1101/2024.01.02.573878

**Authors:** Sebastian Idesis, Michele Allegra, Jakub Vohryzek, Sanz Perl Yonatan, Nicholas V. Metcalf, Joseph C. Griffis, Maurizio Corbetta, Gordon L. Shulman, Gustavo Deco

**Affiliations:** Center for Brain and Cognition (CBC), Department of Information Technologies and Communications (DTIC), Pompeu Fabra University, Edifici Mercè Rodoreda, Carrer Trias i Fargas 25-27, 08005, Barcelona, Catalonia, Spain; Padova Neuroscience Center (PNC), University of Padova, via Orus 2/B, 35129, Padova, Italy; Department of Physics and Astronomy “G. Galilei”, University of Padova, via Marzolo 8, 35131, Padova, Italy; Department of Neuroscience (DNS), University of Padova, via Giustiniani 2, 35128, Padova, Italy; Department of Neurology, Washington University School of Medicine, 660 S. Euclid Ave, St. Louis, MO, 63110; Department of Radiology, Washington University School of Medicine, 660 S. Euclid Ave, St. Louis, MO, 63110; VIMM, Venetian Institute of Molecular Medicine (VIMM), Biomedical Foundation, via Orus 2, 35129, Padova, Italy; Institució Catalana de Recerca I Estudis Avançats (ICREA), Passeig Lluis Companys 23, 08010, Barcelona, Catalonia, Spain; Centre for Eudaimonia and Human Flourishing, Linacre College, University of Oxford, UK; Universidad de San Andrés, Buenos Aires, Argentina; National Scientific and Technical Research Council, Buenos Aires, Argentina; Institut du Cerveau et de la Moelle épinière, ICM, Paris, France

**Keywords:** Whole-Brain models, Predictive, Stroke, fMRI, Dynamics

## Abstract

Computational whole-brain models describe the resting activity of each brain region based on a local model, inter-regional functional interactions, and a structural connectome that specifies the strength of inter-regional connections. Strokes damage the healthy structural connectome that forms the backbone of these models and produce large alterations in inter-regional functional interactions. These interactions are typically measured by correlating the timeseries of activity between two brain regions, so-called resting functional connectivity. We show that adding information about the structural disconnections produced by a patient’s lesion to a whole-brain model previously trained on structural and functional data from a large cohort of healthy subjects predicts the resting functional connectivity of the patient about as well as fitting the model directly to the patient’s data. Furthermore, the model dynamics reproduce functional connectivity-based measures that are typically abnormal in stroke patients as well as measures that specifically isolate these abnormalities. Therefore, although whole-brain models typically involve a large number of free parameters, the results show that even after fixing those parameters, the model reproduces results from a population very different than the population on which the model was trained. In addition to validating the model, these results show that the model mechanistically captures relationships between the anatomical structure and functional activity of the human brain.

## 1. Introduction

A significant development in neuroscience has involved the conceptualization of the brain as a set of dynamic networks that interact and facilitate information processing through the integration and segregation of information. Correspondingly, the application of formal methods from graph theory ^1,2^ and statistical mechanics for studying the structure and dynamics of those networks ^3^ has been essential to this development. The spatiotemporal activity of the brain’s resting-state physiology is identified by measuring the inter-regional correlation of the BOLD signal ^4^, so-called resting-state functional connectivity (FC). This spatiotemporal activity arises and is constrained by a structural connectome ^5^, a small-world organization ^6^ in which ‘wiring efficiency’ is maximized by groups of densely interacting sets of brain regions, i.e. networks, that are linked by sparser connections. The structural organization of the brain generates spatiotemporal dynamics of activity, or recurring waves across cortical, subcortical, and cerebellar circuits ^7^, which occur within a critical regime ^8,9^.

The last decade has seen the development of computational models that simulate the spatiotemporal patterns of brain activity by combining biologically plausible whole-brain descriptions of the macroscale structural connectivity with models of local regional activity. These “whole-brain models” have been able to replicate at least at the group level, many of the spatial and temporal properties of brain activity ^10^. Whole-brain models have also simulated activity changes produced by behavioral states, drugs, or neurostimulation in the healthy or pathological brain ^11,12^.

A key health-related application of whole-brain models has been the simulation of the effects of brain pathologies such as stroke, epilepsy, or schizophrenia on brain activity and behavior ^13,14^. Pathology-specific whole-brain models are very important since accurate simulations of the physiological effects of a pathology and their distributed impact on brain networks may not only provide insights into disease mechanisms, but may additionally allow the effect of therapeutic interventions such as drugs, rehabilitation or stimulation to be modeled or predicted ^11^.

Our group has pioneered the study of brain network alterations in stroke, and their simulation in whole-brain models. Focal lesions like stroke produce characteristic patterns of behavioral impairment and alterations in structural-functional connectivity measured with magnetic resonance imaging ^15–21^. In parallel, we and others have been able to show that whole-brain models can reproduce the abnormalities in network segregation, integration, variability, and criticality of neural states that are observed following a stroke ^13,22–25^.

However, whole-brain computational models often involve large numbers of free parameters, making validation a critical issue, which has been addressed using variants of out-of-sample prediction. In a leave-one-out procedure, the model is fit to all members of the sample except for one, whose data is ‘predicted’. This procedure is then successively applied to each member of the sample. In a cross-validation procedure, the model is applied to a completely different sample from the same training population. However, a more demanding validation criterion would provide better support. Here we fit the model to a sample from one population, i.e. healthy participants, and then apply it to a sample from a completely different population, i.e. stroke patients. ‘Out-of-population’ prediction can be implemented for a stroke population since the effects of a sub-acute stroke lesion on an important measure of whole-brain function, i.e. resting-state FC, are primarily determined by the effect of the lesion on the brain’s structural connectome^17^, and critically, the alterations in a patient’s structural connectome due to a lesion can be incorporated into the whole-brain model without introducing new parameters. Moreover, in addition to validating the model, the application of the out-of-population procedure to stroke patients specifically tests how well the model integrates functional brain activity with structural connectivity, since the predicted physiology in a patient will only be accurate to the extent that the alterations in structural connectivity produced by their lesion appropriately modify the model’s outputs.

We apply this more demanding validation criterion to a whole-brain model that is generative, i.e., that generates BOLD timeseries across a participant’s brain. One advantage of a generative model is that it allows the prediction of any functional brain measure that can be computed from a BOLD timeseries, i.e., the model is not restricted to predicting FC. In addition, the generative aspect of the model allows for generalization to new datasets.

Another noteworthy aspect of the generative, predictive whole-brain model of stroke that is evaluated in this paper is that it predicts measures of brain function and behavior for individual patients. Briefly, a ‘healthy’ model is first derived in a group of healthy control subjects by combining local, parcel-level measures of activity from a Hopf model^26,27^ with a population-level structural connectome^28^ that is modified during model fitting using directional connectivity parameters (GEC parameters) that instantiate generative effective connectivity. The resulting healthy model is then ‘damaged’ separately in each patient by using the structural disconnections produced by their lesion to proportionally modify the GEC parameters, a modification that does introduce any new parameters. Finally, the model-derived activity timeseries for each patient are convolved with a hemodynamic response function to generate blood oxygenation level dependent (BOLD) timeseries across the brain. This generative, predictive whole-brain model, which satisfies an out-of-population validation criterion, differs from our previous stroke-related models since the latter were fit directly to the functional data of each patient.^25^ Consequently, our previous models were not predictive and could not reproduce the functional connectivity anomalies observed in patients from their structural alterations alone.

The results described below validate the new predictive model by showing that it reproduces FC abnormalities, both at the individual and group-level, that resemble the empirical findings reported in the literature. Moreover, the new model has the same degree of accuracy as our previous models, which were fit directly to patient’s functional data. In addition to supporting model validation, these results show that the model mechanistically captures relationships between anatomical structure and functional activity in the human brain.

## 2. Methods

### 2.1 Subjects

We used the Washington University Stroke Cohort dataset ^15^, a large prospective longitudinal (two weeks, three months, 12 months) study of patients with a first-time, single lesion stroke. Although patients were studied at 1-3 weeks (mean = 13.4 days, SD = 4.8 days), 3 months, and 12 months after stroke onset, the current study only analyzed data from the first time point. Furthermore, a group of age-matched control subjects was evaluated twice at an interval of three months. From this cohort, we selected 96 stroke patients and 27 healthy subjects.

Stroke patients were prospectively recruited from the stroke service at Barnes-Jewish Hospital (BJH), with the help of the Washington University Cognitive Rehabilitation Research Group (CRRG). The complete data collection protocol is described in full detail in a previous publication ^15^. Healthy controls were selected based on the same inclusion/exclusion criteria as in ^15^. This group was typically constituted of spouses or first-degree relatives of the patients, and were age- and education-matched to the stroke sample. Patients were characterized with a robust neuroimaging battery for structural and functional features, and an extensive (∼2 hour) neuropsychological battery.

### 2.2 Neuroimaging acquisition and preprocessing

A complete description of the neuroimaging assessment is given in ^17^. Neuroimaging data were collected at the Washington University School of Medicine using a Siemens 3T Tim-Trio scanner with a 12-channel head coil, specifically: 1) sagittal T1-weighted MP-RAGE (TR=1950 msec; TE=2.26 msec, flip angle = 90 degrees; voxel dimensions = 1.0×1.0×1.0 mm), and 2) a gradient echo EPI (TR=2000 msec; TE=2 msec; 32 contiguous slices; 4×4 mm in-plane resolution) resting-state functional MRI scans from each subject. Participants were instructed to fixate on a small centrally located white fixation cross that was presented against a black background on a screen at the back of the magnet bore. Between six and eight resting-state scans (128 volumes each) were obtained from each participant (∼30 minutes total) giving a total of 896 time points for each participant.

Resting-state fMRI preprocessing included (i) regression of head motion parameters, signals from the ventricles, CSF, and white matter, and the global signal (ii) temporal filtering retaining frequencies in the 0.009 - 0.08 Hz band: and (iii) censoring of frames with large head movements, FD = 0.5 mm. The resulting residual time series were projected onto the cortical and subcortical surface of each subject’s brain, which was divided into 234 regions of interest (200 cortical plus 34 subcortical). These regions were taken from the multi-resolution functional connectivity-based cortical parcellations developed by Schaefer and colleagues ^29^, including additional subcortical and cerebellar parcels from the Automated Anatomical Labeling (AAL) atlas ^30^ and a brainstem parcel from the Harvard-Oxford Subcortical atlas (https://fsl.fmrib.ox.ac.uk/fsl/fslwiki/Atlases).

A structural connectome atlas was created using a publicly available diffusion MRI streamline tractography atlas based on high angular resolution diffusion MRI data collected from 842 healthy Human Connectome Project participants ^28^ as described previously (Griffis et al., 2019, 2021). Briefly, the HCP-842 atlas was built using high spatial and high angular resolution diffusion MRI data collected from N=842 healthy Human Connectome Project participants. These data were reconstructed in the MNI template space using q-space diffeomorphic reconstruction ^31^, and the resulting spin distribution functions were averaged across all 842 individuals to estimate the normal population-level diffusion patterns. Whole-brain deterministic tractography was then performed on the population-averaged dataset using multiple turning angle thresholds to obtain 500,000 population-level streamline trajectories.

### 2.3 Neuropsychological and behavioral assessment

The same subjects (controls and patients) underwent a battery of neuropsychological tests at each time point. Briefly, the battery consisted of 44 measures across four domains of function: language, motor, attention, and memory (for a complete description of the tasks measures, see ^15^), as well as a perimetric assessment of visual fields. A dimensionality reduction was applied to the individual test data in each domain using principal component analysis as in ^15^, yielding summary domain scores: Language, MotorR and MotorL (one score per side of the body), AttentionVF (visuospatial field bias), Attention average performance (overall performance and reaction times on the battery’s attention tasks), and AttentionValDis (the ability to reorient attention to unattended stimuli), MemoryV (composite verbal memory score) and MemoryS (composite spatial memory score). Finally, patients’ behavioral scores were z-scored with respect to controls’ scores, to highlight behavioral impairments.

In addition to domain-specific scores, the patients’ clinical severity was assessed through the National Institutes of Health Stroke Scale (NIHSS) ^32^ that includes 15 subtests addressing: level of consciousness (LOC), gaze and visual field deficits, facial palsy, upper and lower motor deficits, limb ataxia, sensory impairment, inattention, dysarthria and language deficits. The total NIHSS score was used as an averaged measure of clinical severity for each patient.

### 2.4 Functional connectivity (FC) measures

Based on previous work ^16,17^ we defined three measures that are consistently impaired in stroke patients:

1. Intra-hemispheric FC: average pairwise FC between regions of the Dorsal Attention Network (DAN) and Default Mode Network (DMN).
2. Inter-hemispheric FC: average homotopic inter-hemispheric connectivity within each network
3. Modularity: overall Newman’s modularity among cortical networks, a comparison between the number of connections within a module to the number of connections between modules ^33^

### 2.5 Lesions

Each lesion was manually segmented on structural MRI scans and checked by two board-certified neurologists. The location (cortico-subcortical, subcortical, white-matter only) of each lesion was assigned with an unsupervised K-means clustering on the percentage of total cortical/subcortical gray and white matter masks overlay. For a more extensive explanation on how the overlap of each lesion group with gray matter, white matter, and subcortical nuclei is calculated, see Corbetta et al, 2015.

### 2.6 Lesion disconnection masks

The Lesion Quantification Toolkit ^34^ produces a comprehensive set of atlas-derived lesion measures that includes measures of grey matter damage, white matter disconnection, and alterations of higher-order brain network topology. Importantly, the measures produced by the toolkit are based on population-scale (e.g., N = 842) atlases of grey matter parcel boundaries and white matter connection trajectories that were constructed from high-quality resting-state functional MRI and diffusion MRI data using state-of-the-art methods ^28^.

Taking advantage of the Lesion Quantification Toolkit (LQT), the structural disconnection (SDC) masks consisted of a spared connection adjacency matrix where each cell quantified the percentage of streamlines connecting each region pair in the atlas-based structural connectome that were spared by the lesion. Therefore, the multiplication of each SDC with the above-mentioned group average SC template ^34^ provides an atlas-based weight for each region pair in each patient. The SDC masks were computed by embedding the lesion in the healthy structural connectivity atlas. Therefore, the measures of structural disconnection were indirectly derived. In the same cohort, we also measured diffusion tensor imaging (DTI) at 3- and 12-months post-stroke, but these timepoints were not included in this study.

Since many stroke lesions occur predominantly in the white matter, or include both a gray and white matter component, the SDC mask should provide an accurate description of the damage to the connectome. We computed the total amount of disconnection ^17^ as a metric of anatomical impairment to assess the validity of the model.

### 2.7 Whole-brain Hopf model parameter estimation

The Hopf model directly simulates BOLD activity at the whole-brain level. The model consists of coupled dynamical units representing the cortical and subcortical brain areas from a given parcellation. The local dynamics of each brain area (node) is described by the normal form of a supercritical Hopf bifurcation, also called a Stuart-Landau Oscillator, which is the canonical model for studying the transition from a fixed point to a limit cycle. We simulated the BOLD activity at the whole-brain level by using the Hopf computational model, which simulates the dynamics emerging from the mutual interactions between brain areas, considered to be interconnected based on the established graphs of anatomical SC ^35,36^. The structural connectivity matrix (group average SC template) was scaled to a maximum value of 0.2 ^36^ in order to explore the range of the G parameter established in previous work. To calculate the Generative Effective Connectivity (GEC), we optimized the phase of the empirically measured FC in the healthy subject group with the phase of the model FC time series. This optimization was carried by changing the global coupling parameter G (obtaining a value of G = 0.75 as the optimal one), which assesses the influence of SC in the model. The higher the value of the factor G, the bigger the influence of the system in each node. The model consists of 234 coupled dynamical units (ROIs or nodes) representing the 200 cortical and 34 subcortical brain areas from the parcellation. When combined with brain network anatomy (explained above in the “Neuroimaging acquisition and preprocessing” section), the complex interactions between Hopf oscillators have been shown to successfully replicate features of brain dynamics observed in fMRI ^35,36^.

In complex coordinates, each node j is described by the following equation: (For more information, see Deco et al., 2019)

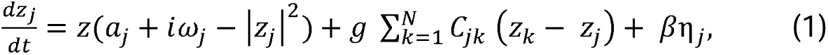

and

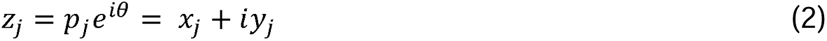

Where *α* and *ω* the bifurcation parameters and the intrinsic frequencies of the system, respectively. This normal form has a supercritical bifurcation at *a_j_*□=□0 for which we used the homogeneous parameter space around the Hopf bifurcation (a = −0.01). Within this model, the intrinsic frequency *ω_j_* of each node is in the 0.04–0.07Hz band (j=1, …, n). The intrinsic frequencies were estimated from the data, as given by the averaged peak frequency of the narrowband BOLD signals of each brain region. The variable G represents a global coupling factor scaling the structural connectivity C_jk_, and η is a Gaussian noise vector with standard deviation *β* = 0.04. This model can be interpreted as an extension of the Kuramoto model with amplitude variations, hence the choice of coupling (*z_k_* - z*_j_*), which relates to a tendency of synchronization between two coupled nodes. We insert equation 2 in equation 1 and separate real part in equation 3 and imaginary part in equation 4 ^36^.

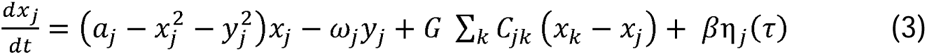

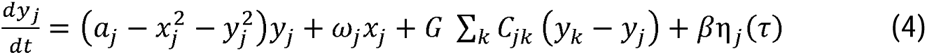

For all cases, we will compute the goodness of fit by the mean error (squared difference) between the upper triangular values of the empirical and simulated FC.

### 2.8 Generative Effective Connectivity calculation

Generative Effective connectivity (GEC) utilizes differences detected at different times in the signals for connected pair of brain regions to infer what effects one brain region has on the other.

The analysis of GEC incorporates an indirect metric (as it is derived from other presented metrics) into the whole-brain model to replace the existing descriptive metrics of FC and SC. Previous studies have shown how GEC is fundamental for understanding the propagation of information in structural networks ^37,38^. Methods for estimating GEC are explained in detail in a previous publication ^39^. Briefly, we computed the distance between our model and the empirical grand average phase coherence matrices (as a measure of synchronization of the system) of the healthy control group and adjusted each structural connection separately using a greedy version of the gradient-descent approach. To work only positive values for the algorithm, all values are transformed into a mutual information measure (assuming Gaussian distribution). We obtained the healthy simulated functional connectivity *FC^model^* from the first N rows and columns of the covariance *K*, which corresponds to the real part of the dynamics (precisely representing the BOLD fMRI signal). We then fit “C” such that the model optimally reproduces the empirically measured covariances *FC^empirical^* (i.e., the normalized covariance matrix of the functional neuroimaging data) and the empirical time-shifted covariances *FS^empirical^* (*τ*) where *τ* is the time lag, which are normalized for each pair of regions *i* and *j* by 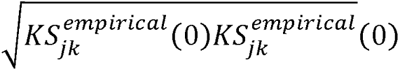. We proceeded to update the C until the fit is fully optimized. The equation of the optimization is as follows: (For more information, see ^40^)

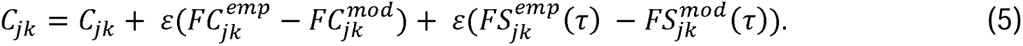

Where C is the anatomical connectivity and is updated with the difference between the grand-averaged phase coherence matrices (Empirical: 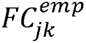 and model: 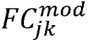) and the difference between the time-shifted covariance matrices, both scaled by a factor *ε* = 0.001. Where 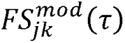 is defined similarly to 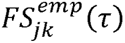. After this process, C is considered as a Generative Effective Connectivity (GEC) matrix. The prediction, therefore, is based on the current estimation of the healthy structural connectivity, which gets updated optimizing the phase FC in each iteration. In summary, the model was run repeatedly with recursive updates of GEC until convergence was reached. The distinction between functional and effective connectivity is crucial here: FC is defined as the statistical dependence between distant neurophysiological activities, whereas GEC is defined as the influence one neural system exerts over another providing directionality in the relations making the matrices asymmetrical ^41,42^.

### 2.9 Models

#### 2.9.1 Full Predictive model

We calculated a predictive model to capture the dynamical effects of stroke lesions two weeks after onset. First, we estimated the optimal value of global coupling for which the modeled Hilbert phases were most similar to the empirical data in the healthy controls group (G=0.75). Then, we computed the GEC (See previous section) on the same group **(Figure 1a)**. Lastly, we added the information of the SDC mask of each patient to the existing GEC to simulate fMRI BOLD data of the corresponding patient **(Figure 1b)**. As a result, the simulated time series (referred to as “Full Predictive model”) contains structural information from the patient but not their functional information, making it a predictive model, in contrast to the previously discussed non-predictive models.

**Figure 1:**
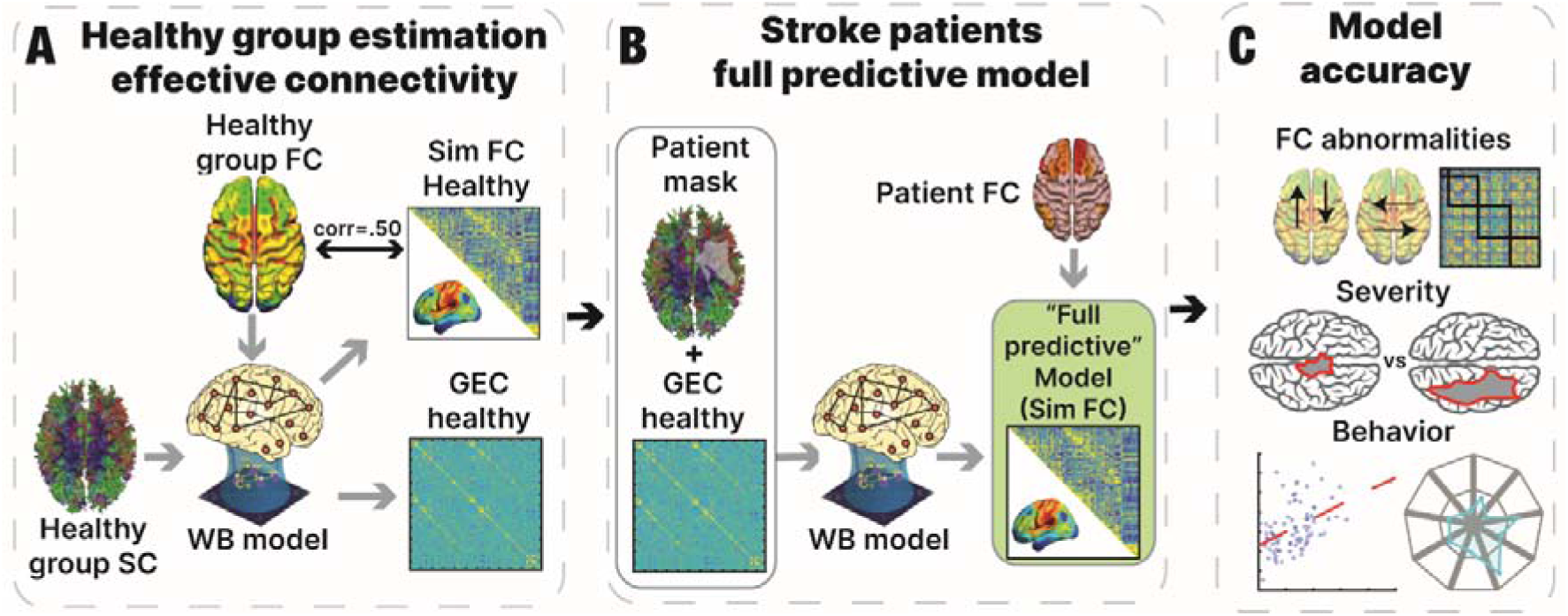
Pipeline for the predictive model: **(A)** Healthy control generative effective connectivity (GEC) was calculated by using the healthy template SC with each healthy control fMRI time series. The model was optimized using a whole-brain (WB) model to create an average GEC for the healthy controls. **(B)** The predictive model used each patient’s disconnection mask to modify the control GEC and obtain the patient’s simulated FC, referred to in the figure as the full predictive model. **(C)** We determined the accuracy of the healthy model in accounting for the FC matrix of each healthy control subject **(Figure 2A)**. We also determined the accuracy of the Full Predictive model in predicting each patient’ FC matrix **(Figure 2A)**, FC-derived measures in each patient that are typically abnormal following a stroke **(Figure 2B)**, z-scored abnormalities, relative to healthy controls, in the FC matrix of each patient **(Figure 2C)**, and each patient’s behavioral deficits (**Figure 2D**, upper panel). We also investigated the determinants of model accuracy by examining whether the accuracy of a patient’s simulated FC matrix covaried with the severity of their lesion-induced structural damage (**Figure 3A**), the magnitude of FC-based and graph-related functional measures that are typically abnormal following a stroke **(Figures 3B-3E)**, and the magnitude of the patient’s behavioral deficits **(Figure 3F)**.

For each patient, the simulated fMRI BOLD timeseries for each parcel pair were then correlated to construct the patient’s simulated FC matrix. To isolate the degree to which the patient’s FC between two parcels was abnormal relative to healthy controls, empirical and simulated FC matrices for each patient were z-scored with respect to the healthy controls’ empirical and simulated FC matrices to create the patient’s empirical and simulated z-scored FC abnormality matrices. Specifically, the healthy group-mean FC for a parcel pair was subtracted from the patient’s FC for that parcel pair, and this difference score was then divided by the standard deviation of the healthy group FC for that parcel pair.

In addition, we separately averaged the homotopic interhemispheric FC entries and the DAN-DMN entries from a patient’s z-scored FC abnormality matrix to create averaged abnormality scores for these two classes of FC, which are typically abnormal in patients.

#### 2.9.2 Predictive comparative models

Three different predictive models were calculated to compare their performance with the full predictive model:

- Predictive model without mask: To assess the effect of incorporating a disconnection mask (lesion information) in the predictive model, this model simply consisted of the healthy group model without any disconnection mask.
- Surrogate mask model: As the effect of the disconnection mask could simply reflect the overall magnitude of disconnection, we computed predictive models in which each patient received the disconnection mask of another patient. As the lesion severity averages out across patients but the pattern/location of the lesion is different, comparisons of the full predictive model vs. the surrogate mask model indicate how strongly the accuracy of a predictive model depends on incorporating the specific features of a patient’s lesion. Therefore, the model without mask and the surrogate mask models serve as predictive controls for the full predictive model.
- G-DSC model: In order to estimate the contribution of the GEC parameters to the accuracy of the Full Predictive model, we constructed a healthy model that was fit using only the G parameter, not the GEC parameters. Therefore, the connections between parcels in this healthy model were based on a healthy structural connectome ^28^ that was not modified via the GEC parameters. As for the Full Predictive model, the healthy model was then lesioned separately for each patient using the patient’s structural disconnection matrix.

#### 2.9.3 “Non-predictive and patient-specific” Model

A non-predictive patient-specific model was calculated for comparison with the Full Predictive model. The patient-specific model used each patient’s own functional information (i.e., their BOLD timeseries) as well as their structural disconnection matrix to estimate the parameters of their model, making the model non-predictive.

### 2.10 Assessment of model accuracy

#### 2.10.1 Behavior impairment prediction

We explored how well subjects’ behavioral scores (See section 2.3) were predicted by their empirical and simulated FC. We calculated two partial least-squares regression (PLSR) models using the empirical and simulated FC matrices as predictors. As a control, we included a third model based solely on anatomical information (the SDC matrix), as reported in previous literature ^16,17,43^.

PLSR is a multivariate regression technique ^44^ that is closely related to principal components regression ^45^. Both approaches are especially useful for situations where there are more variables than observations and/or when there is high collinearity among the predictor variables. Nevertheless, PLSR has important advantages that are primarily due to differences in the criteria used for decomposition of the predictor matrix ^17^. Detailed descriptions of theory and algorithms behind the PLSR approach are explained in previous literature ^46^

#### 2.10.2 Global efficiency

Global efficiency was calculated as the average inverse shortest path length ^47^. Unlike path length, the global efficiency can be calculated on disconnected networks, as paths between disconnected nodes are defined to have infinite length, and correspondingly zero efficiency. Therefore, it is an ideal metric when investigating stroke data. In contrast to path length, which is primarily influenced by long paths, global efficiency is primarily influenced by short paths. Different authors have claimed that this may convert global efficiency into a superior metric of integration ^48,49^. Global efficiency is calculated as follows ^47,49^:

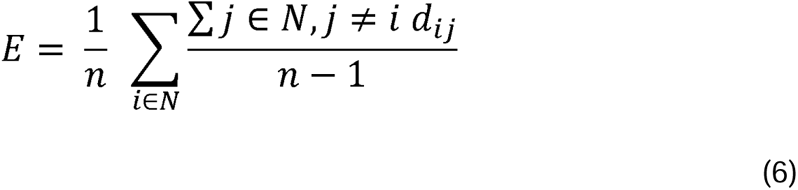

Where N is the set of all nodes in the network, n is the number of nodes, and (*ij*) is a link between the nodes i and j.

#### 2.10.3 FC Entropy

FC entropy is an information theoretical metric that measures the richness of functional connections and therefore may be a relevant biomarker for many disorders^24,25,50^.

Previous studies have reported abnormal FC entropy values when comparing healthy controls with stroke patients ^22,24^. However, these models use generic anatomical connectomes based on group averages instead of personalized structural connectivity. Although the current study used an atlas-based structural connectome for modeling the healthy control subjects, this connectome was separately modified for each patient based on their lesion.

Entropy is calculated as follows ^25^:

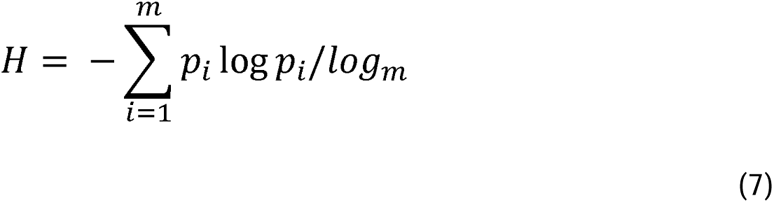

Where m is the number of bins used to construct the probability distribution function of the upper triangular elements of |FC|. The normalization factor in the denominator is the entropy of a uniform distribution, and it ensures that H is normalized between 0 and 1.

#### 2.10.4 Average degree

Average degree is a measure of overall network connectivity that provides information about network segregation and integration ^49^.

Average degree is calculated as follows ^25^:

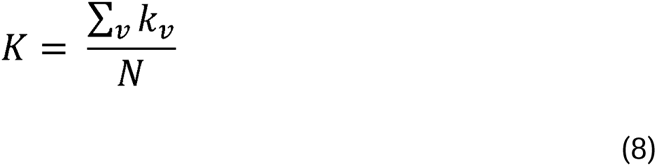

Where N is the number of nodes and *k_v_* is the degree of the node v as defined above.

## 3. Results

To infer the dynamical effects of stroke lesions two weeks after onset, we used a computational model based on coupled Stuart Landau oscillators (**Figure 1A**). The model contains a global scale factor, also referred to as the G coupling value, which determines the influence of SC in the model. It also contains Generative effective connectivity (GEC) parameters that capture directional interactions between regions. Both types of parameters are optimized to improve the model fit (i.e., the similarity of empirical and model FC). In the current study, we use only the functional data of the healthy control dataset to optimize these parameters at the group level. Performing an exhaustive exploration of the homogeneous parameter space (a, G) around the Hopf bifurcation (a = −0.01), we found G = 0.75 as the optimal value of G for which the modeled FC of the Hilbert phases were most similar to those observed in the empirical data. Initializing the GEC to be equal to the SC, we iteratively adjusted its values to improve the similarity between the model and empirical FC at a group level. **Supp. Figure 1** shows the role of the structural connections between parcels by assessing the difference between the empirical and simulated FC. We then added information about the structural damage caused by a patient’s stroke to the healthy group GEC by using a structural disconnection mask to create a predictive model that generated a simulated version of each patient’s FC matrix (**Figure 1B**). Therefore, the Full Predictive model reproduces the functional consequences of stroke lesions in individual patients by exploiting the patient’s structural disconnection matrix. Importantly, in contrast to previously reported models, the Full Predictive model uses the functional data of only healthy control subjects to predict a patient’s FC.

From the obtained simulations, several metrics were calculated to assess how well the predictive model captured the functional effects of stroke. For this purpose, among other results, we considered the main FC-based metrics that are known to give abnormal values in stroke patients (mean intra-hemispheric FC between the DAN and DMN, mean homotopic interhemispheric FC, and modularity level). Similarly, we determined how well the predictive model captured patients’ behavioral deficits. To better understand the determinants of model accuracy, we also examined whether the accuracy of the predictive model for a patient covaried with the magnitude of structural, FC-based and graph-based measures that index the severity of the patient’s stroke, such as total structural disconnection, modularity, and global efficiency (**Figure 1C**).

### 3.1 Model outcomes and their relationship with stroke effects

We first determined how well the model accounted for each subject’s empirical FC matrix (healthy controls and stroke patients) by computing the Pearson correlation between the model-generated FC matrix and their empirical FC matrix. **Figure 2A** shows the distribution of correlation coefficients over the sample, indicating that both the healthy model and the Full Predictive model generated simulated FC matrices that showed a moderate level of accuracy (healthy group mean, r=0.43, std=.05; patient group mean, r=0.37, std=.05; **Figure 2A**).

**Figure 2:**
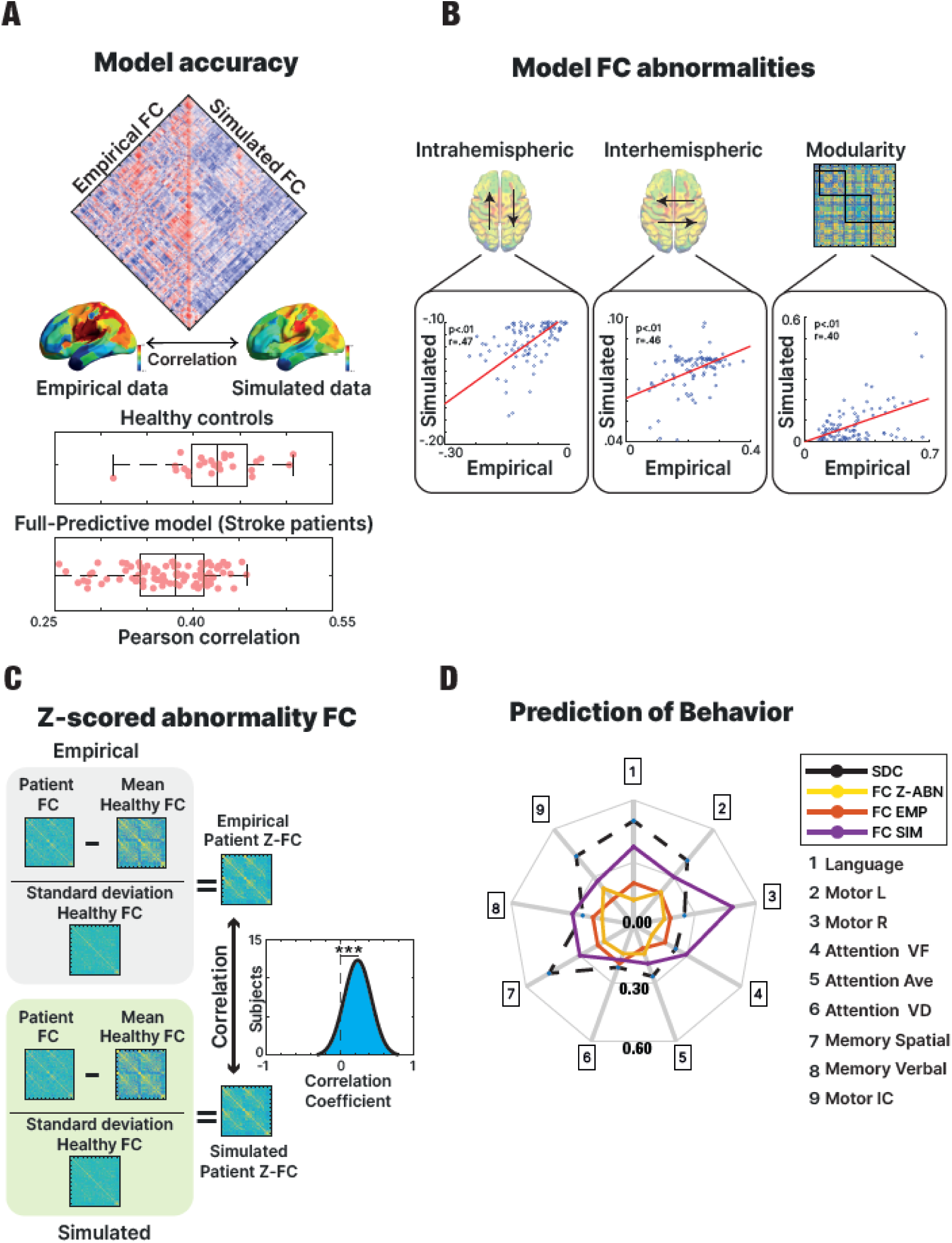
Model prediction of patient FC and behavior. **(A)** The Pearson correlation between the empirical and simulated FC matrices of each subject (healthy controls and stroke patients) was computed in order to assess the accuracy of the models. **(B)** FC-based measures typically abnormal in stroke patients were calculated from the empirical and simulated FC matrices from the Full Predictive model in order to compare their similarity. **(C)** The distribution across patients of the correlation between the empirical and simulated z-scored FC abnormality matrices for each patient from the Full Predictive model. **(D)** Separate partial least squares regression (PLSR) analyses were conducted using the empirical, z-scored and simulated FC matrices from the Full Predictive model and the structural disconnection matrix as regressors to predict each domain of behavioral impairment.

To assess whether the Full Predictive model accurately predicted specifically the abnormalities in a patient’s FC that resulted from their stroke, we correlated each patient’s empirical and simulated z-scored FC abnormality matrices. **Figure 2C** shows that the mean correlation across patients (*r*=.24, *std*=.18) was significantly positive *(t*(95)= 12.65, *p*< .01), indicating that patient abnormalities in FC were significantly predicted.

Next, we analyzed how well the model predicted specific FC-based measures that are typically abnormal in stroke patients: 1) a decrease of negative intra-hemispheric FC between regions of the Dorsal Attention Networks (DAN) and Default Mode Network (DMN); 2) a decrease of inter-hemispheric homotopic FC; 3) a decrease of modularity. **Figure 2B** shows that the empirical and simulated values for these three signatures of stroke were significantly correlated across patients (intra-hemispheric: *r*=.47, *p*<.01; inter-hemispheric: *r*=.46, *p*<.01; modularity: *r*=.40, *p*=.03). Previous studies have shown that whole-brain GEC models preserve the same three FC-based measures, especially when they include structural disconnection information, revealing the key importance of incorporating this information into the models ^13^. In addition, for each patient we separately averaged the entries in their z-scored FC abnormality matrix for interhemispheric homotopic FC and DAN-DMN FC to assess whether the model specifically predicted patient abnormalities in these two FC measures. The results, shown in **Supplementary Figure 2**, indicate that abnormalities in both FC measures were significantly predicted. Overall, the above results indicate that the Full Predictive model reproduced to some extent patient FC matrices, patient-specific abnormalities in the FC matrix, summary FC-based measures that are typically abnormal following a stroke, and patient-specific abnormalities in those measures.

Finally, we used Partial Least Squares Regression (see methods) to separately assess how well the simulated FC matrices, the empirical FC matrix, the z-scored FC abnormality matrix, and the SDC matrix predicted the patients’ behavioral scores. Although the simulated FC matrix was modestly predictive across behavioral domains, the model independent SDC matrix showed nominally better performance except for the domains for Attention VF, Attention Ave and Memory Verbal (**Figure 2D**). The empirical FC matrix showed lower performance than the z-scored and simulated FC matrices across all domains, possibly because the predictive model that generated a patient’s simulated FC matrix explicitly incorporated their structural disconnection matrix. Overall, these results indicate that although the predictive model partly accounted for behavioral abnormalities, the model did not generally perform better than a purely structural measure.

### 3.2 Model accuracy in relation to structural damage and global metrics

Next, we investigated whether structural and functional features influenced how accurately the model accounts for the data from individual patients. Specifically, we examined whether the accuracy of the model’s simulation of a patient’s FC matrix, as indexed by the correlation between the patient’s simulated and empirical FC matrices, covaried with the structural damage from the patient’s own lesion, the values of the patient’s graph-based functional metrics, and the magnitude of FC abnormality metrics that are canonically affected in stroke.

We found that higher model accuracy was associated with lower values of total structural disconnection (R: −.58, p<.01), which served as a measure of overall lesion damage (**Figure 3A, left panel**). When splitting the sample in half by using the median value of total structural disconnection, patients with greater total disconnection showed significantly lower model accuracy (*t*(94)= 5.82, *p*< .01) (**Figure 3A, right panel**). A similar analysis using lesion volume (number of damaged voxels) as an alternative metric yielded similar results (**Supp Figure 3**; an example of a patient lesion and the corresponding asymmetric effective connectivity matrix is shown in **Supp. Figure 4)**.

**Figure 3:**
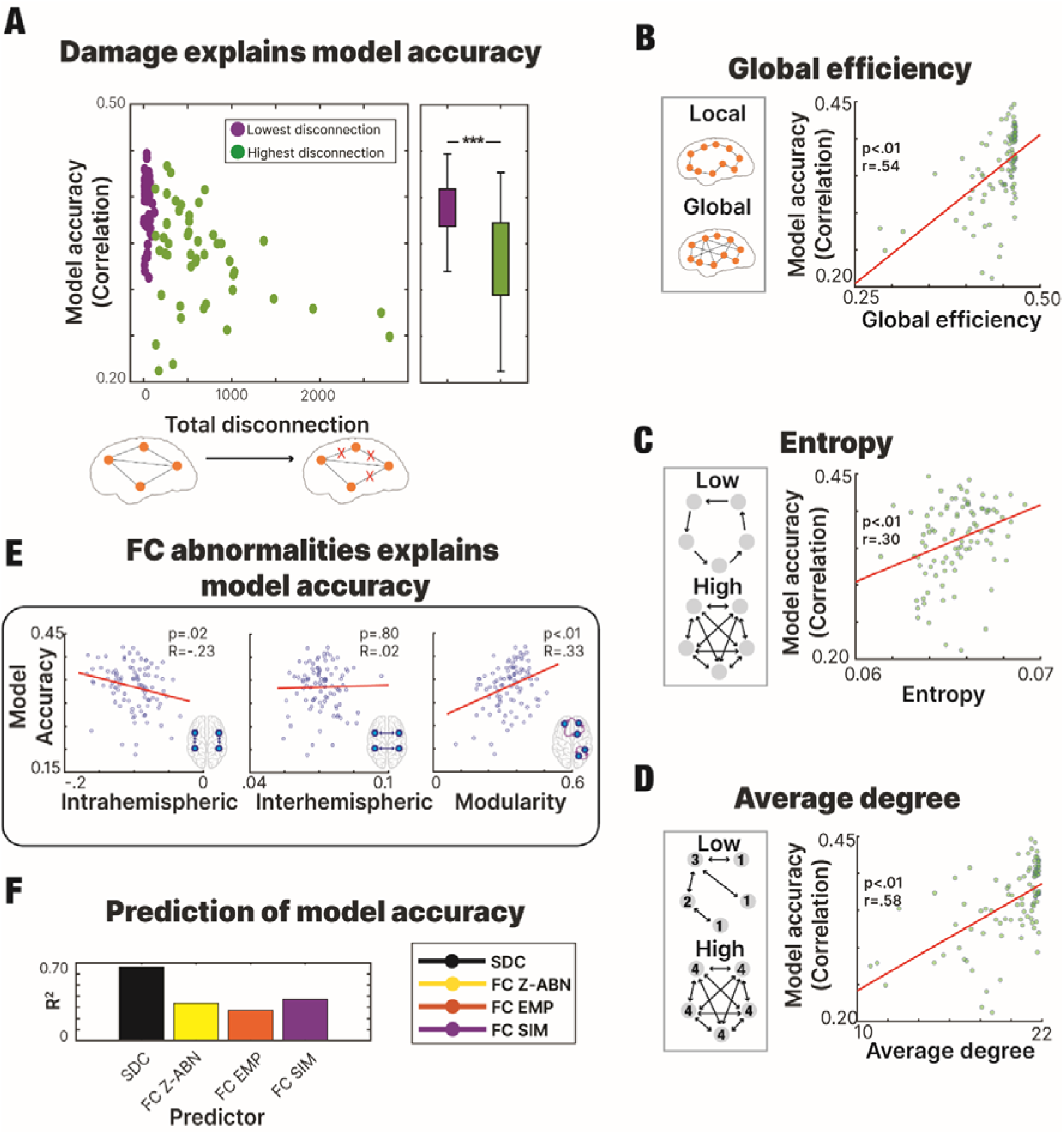
Structural and functional determinants of model accuracy for individual patients **(A)** Subjects with lower levels of disconnection exhibited a higher correlation between the empirical and simulated FC matrices, indicating better model performance for patients with less severe lesions. **(B-C-D)** Higher global efficiency **(B)**, entropy **(C)** and average degree **(D)** were associated with higher model accuracy. **(E)** Model accuracy for FC-based measures typically abnormal in stroke patients was assessed for the presented model. Accuracy of the full predictive model was significantly associated with the magnitude of intrahemispheric FC and modularity but not interhemispheric FC. **(F)** Model accuracy predicted by each type of regressor (SDC matrix, FC empirical and FC simulated matrices, z-scored FC abnormality matrix) in a PLSR analysis of model accuracy.

The dependence of model accuracy on lesion severity was consistent with its dependence on the magnitude of graph-based metrics that are typically abnormal following a stroke ^25^. We found significant positive correlations between model accuracy and global efficiency (*r*=.54, *p*<.01; **Figure 3B**), entropy (*r*=.30, *p*<.01; **Figure 3C**), and average degree (*r*=.58, *p*<.01; **Figure 3D**). In other words, model accuracy was higher when the lesion produced weaker network function abnormalities.

Furthermore, the Full Predictive whole brain model generated FC matrices whose correlation with empirical FC matrices (i.e., model accuracy) was significantly related to the magnitudes of intra-hemispheric FC (*R*=-.23, *p*=.02) and FC modularity (*R*=.33, *p*<.01) as seen in **Figure 3E**. The sign of the relationship was consistent with the conclusion from **Figures 3A** and **3B-D** that the model more poorly predicted the functional measures of patients that had more abnormal structural or functional measures. However, model accuracy was not correlated with the magnitude of inter-hemispheric FC, which is typically lower in stroke patients than controls.

Finally, separate PLSR analyses showed that model accuracy for a patient was well predicted by the full SDC matrix, next by the simulated FC matrix, the z-scored FC matrix and least by the empirical FC matrix (SDC: R^2^=0.66, p<.01, Sim-FC: R^2^: 0.37, p<.01, Z-scored Abnormality FC: R^2^: 0.35, p<.01, and Emp-FC: R^2^: 0.27, p<.01) (**Figure 3F**).

Overall, these results indicate that the model’s ability to accurately reproduce a patient’s FC matrix decreased as the patient’s structural measures, and to a lesser extent functional measures, showed larger departures from those for healthy controls.

### 3.3 Model Comparisons

Having established the accuracy with which the Full Predictive model reproduced measures that are typically abnormal in stroke patients (**Figures 2B and 2C**), we compared its performance with a patient-specific non-predictive model reported in a previous study ^13^ that used both anatomical and functional information to simulate a patient’s time series (referred to as the Non-predictive patient specific model) (**Figure 4A**).

**Figure. 4:**
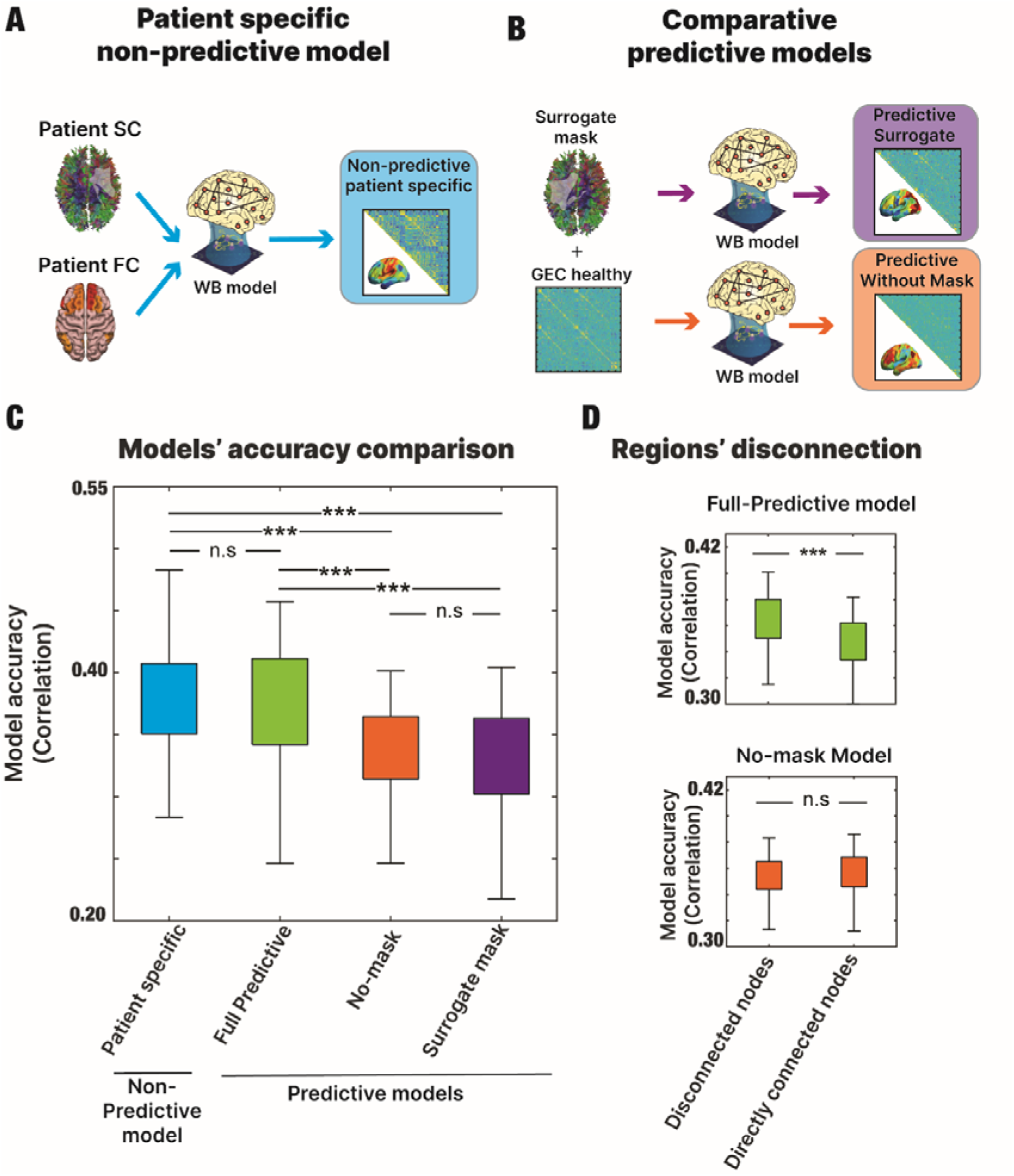
Model comparisons: **(A)** We calculated the Non-predictive patient specific model using both the anatomical and functional data of each patient. This model was not predictive since it was fit to the patient’s functional data. **(B)** We calculated two comparative predictive models to compare with the Full Predictive model. The Predictive No-mask model was built using only the healthy GEC while the predictive surrogate mask model was calculated by modifying the healthy GEC via the disconnection mask of a different patient. **(C)** The similarity between the empirical and simulated FC matrices was assessed for each model. The non-predictive patient specific model and full predictive model showed similar levels of performance that exceeded performance for the surrogate and no-mask models. **(D)** Comparison of model performance when dividing by disconnected nodes and directly connected nodes showed a significant interaction between model and node connection.

Secondly, two comparative predictive models were chosen to assess how much model accuracy depended on the patient’s disconnection mask that was incorporated in the Full Predictive model. The model without mask allowed us to assess the accuracy gain obtained by incorporating the patient’s lesion information, with respect to not incorporating any lesion information (i.e., using the unmodified healthy group model for all patients). The surrogate mask model allowed us to assess the accuracy gain obtained by incorporating specifically the patient’s lesion information, with respect to using a lesion from a different patient (**Figure 4B**).

We compared the performance of all models by computing the correlation of the simulated and empirical FC matrices. The Full Predictive model showed roughly equivalent accuracy to the Non-predictive patient specific model ^13^, while the Model with surrogate mask and the model without mask showed lower accuracy (**Figure 4C**).

A within-subject ANOVA indicated that the main effect of model type (non-predictive, full predictive, no mask, surrogate mask) on accuracy was significant (*F(3,*285)= 14.84, *p*<.01). Post-hoc tests indicated that the patient-specific and full predictive models did not significantly differ in accuracy (p<.36) but were significantly more accurate than both the surrogate mask and no mask models (p<.01 in all cases).

In order to assess the effects of the GEC parameters on model accuracy (see Methods, section 2.9.2, Predictive comparative models, G-DSC model), a healthy model was fit using only the G parameter, not the GEC parameters, and was then lesioned as in the Full Predictive model in order to generate predictions for the patients. The results, shown in **Supplementary Figure 10**, indicate that model accuracy was significantly higher for the Full Predictive model than for the G-DSC model.

Because the accuracy of the Full Predictive model depends on the integration of functional and structural information, we assessed this relationship in more detail. Specifically, model accuracy was computed as a function of whether two parcels were directly or indirectly connected (**Figure 4D**). A two-way ANOVA was performed to analyze the effect of model type and node connection on the model accuracy. The two-way ANOVA revealed that there was a statistically significant interaction (F(1, 190) = 67.96, p < .01). The Full Predictive model showed significantly greater accuracy for indirectly than directly connected parcels (*p*<.01), but no difference was found for the No Mask model (*p>*.05).

Finally, model accuracy relative to the accuracy of the healthy control group model is shown in **Supplementary Figure 5**. The influence of the global coupling parameter (GC) is presented in **Supplementary Figure 6**. The relation between the accuracy of each model and the magnitude of FC measures typically abnormal in stroke patients is presented in **Supplementary Figure 7**. Comparisons of dynamical metrics ^21,51^ between the models are presented in **Supplementary Figure 8** (see figure caption for explanation of metrics).

Overall, the results show the efficacy of the Full Predictive model, which does not use a patient’s functional BOLD data, allowing its predictions to be generalized to new patient datasets, and opening the door for predicting the expected effects of a simulated lesion or external stimulation of a patient’s brain.

## 4. Discussion

The results show that functional connectivity in patients could be predicted by a whole-brain computational model strictly from the structural disconnection caused by a patient’s lesion, suggesting that the model mechanistically captured to some degree the relationship between anatomical structure and functional activity. Moreover, the model significantly predicted abnormalities in patient FC with respect to the FC of the healthy control group Although the model also predicted the behavioral abnormalities of patients, prediction was no better than that obtained using a purely structural measure, the structural disconnection matrix. While previous work has examined how well computational models can reproduce FC when model parameters are directly fit using functional and structural data from healthy controls or stroke patients ^13,52^, the current study moves fundamentally beyond such work by determining whether these models can in fact predict the effects of a stroke based solely on the structural information associated with a patient’s lesion ^25^.

### Validating the Full Predictive model’s integration of structure and function

The introduction noted that the large number of free parameters in whole-brain models emphasizes the need for strong validation procedures, such as the out-of-population approach taken here. Both the fact that the Full Predictive model significantly predicted patient abnormalities and that the accuracy of the predictive model was essentially equivalent to the accuracy obtained by fitting a non-predictive model directly to the patient’s functional and structural data, despite the absence of free parameters in the Full Predictive model, provides support for model validity. Moreover, because the out-of-population prediction was applied to a patient population with a different structural connectome than the training population, the variation of model accuracy with lesion severity, connection type (direct, indirect), and lesion mask type (the patient’s own mask, a different patient’s mask, no mask) provided insights into the conditions under which the Full Predictive model best integrated anatomical structure with functional activity.

We first fit a model to data from age- and education-matched healthy controls based on a healthy structural connectome and the healthy controls’ functional imaging data. For each patient, we then determined how the patient’s lesion has modified the healthy structural connectome and made corresponding changes to the structural connectivity parameters in the healthy model (the GEC parameters) without additional model fitting or recourse to the patient’s functional data. Finally, the modified healthy model specific to the patient, i.e., the Full Predictive mdoel, generated the patient’s predicted FC, which was compared against the empirically measured FC. Specifically, model accuracy was evaluated by assessing the correlation between the patient’s empirical and predicted FC matrices. Because FC matrices specify the functional interactions between each pair of brain regions, the predicted matrices potentially provide information concerning which functional connections are particularly vulnerable in the patient, a possibility also raised by the prediction of patients’ z-scored FC abnormality matrices.

Interestingly, the accuracy of the full predictive model was essentially equivalent to the accuracy obtained by fitting the model directly to the patient’s functional and structural data. Moreover, the accuracy of the Full Predictive model depended on incorporating the structural disconnection specific to that patient’s lesion, as shown by the significantly poorer performance obtained by substituting the structural disconnection for a different patient. Both results provide additional support for how the Full Predictive model integrated structure and function.

However, the accuracy for predicting a patient’s FC matrix tended to be less the more a patient’s structural connectome and functional measures differed from the structural connectome and functional measures of healthy control subjects. Specifically, accuracy decreased with the magnitude of the total structural disconnection caused by the lesion. Similarly, measures of modularity and intra-hemispheric FC and graph-based measures that are typically abnormal following a stroke tended to be more poorly predicted the more they differed from healthy control values (surprisingly, a similar relationship was not observed for inter-hemispheric FC, an important signature of stroke-induced dysfunction).

One interpretation is that the model tended to better predict patients that were more similar to healthy controls since it was initially based on a model computed from healthy control data.

This interpretation suggests a limitation on how well the Full Predictive model integrated anatomical structure and functional activity, an interpretation also suggested by the difference in model accuracy for node pairs that were directly vs. indirectly connected in the healthy structural connectome.

Alternatively, similar relationships may also be present for the non-predictive patient-specific model, i.e., the dependence of model accuracy on structural and functional measures may not be related to prediction per se or be a unique feature of the Full Predictive model. The results in **Supplementary Figures 7 and 9** provide some support for this alternative, but those results do not rule out the first explanation as a contributing factor.

Finally, as a further alternative, it can be considered that the more damage occurred in the brain, the more functional effects appear in secondary and tertiary connections, which are not well estimated in the healthy controls (given that healthy control FC does not change its weights). This reasoning would explain why indirect FC measures are not significantly predictive of behavior while direct FC measures are, especially for cognitive deficits that involve multi-network pattern abnormalities ^16,19^.

### Relation to previous work

Previous computational work based on concepts from statistical mechanics has shown that resting-state organization conforms to a state of ‘criticality’ that promotes responsiveness to external stimulation, i.e. resting state organization facilitates task-based processing ^12,53–55^. The rich body of empirical work on resting-state organization has facilitated an important testing ground for evaluating computational whole-brain models. In these models, neural modules or elements are connected by ‘structural’ links that mirror the empirical structural connectivity of the human brain as assessed using diffusion-based MRI ^27,53,54,56^, resulting in resting-state dynamics that respect critically. Initial applications of whole-brain computational models to stroke populations ^22,24^ used the biophysically-based model of Deco et al. ^54^, which involves a mean field approximation of populations of spiking neurons with realistic NMDA, AMPA, and GABA synaptic dynamics. However, the authors subsequently developed the mesoscopic Hopf model ^39^ used in the current study, which provides a better fit to healthy control data and runs two orders of magnitude faster, allowing the use of higher-resolution functional parcellations that likely increase model accuracy.

The whole-brain computational models presented in recent studies that involve stroke patients included a global coupling parameter and GEC parameters that encoded directional interactions between nodes that had direct structural connections ^13,52^. The resulting generative effective structural connectivity weights allowed a better fit between the empirical and modeled FC than that achieved by models that only varied the global coupling parameter. In both papers, however, the model was fit directly to a patient’s functional data, and therefore was not a predictive model.

The Full Predictive model could be applied in future work to other focal and non-focal pathologies that damage the structural connectome. Although the current study focused on predicting FC in sub-acute stroke patients, future studies could examine whether changes in structural connectivity during recovery also produce predicted changes in FC. Because the dataset consisted mostly of ischemic patients, however, model predictions will need to be tested in hemorrhagic stroke patients before concluding that the model applies more generally to stroke.

### Limitations

The mesoscopic Hopf model ^39^ includes global coupling and GEC parameters that affect the connectivity between nodes of the model and bifurcation parameters that affect the dynamics of the nodes (the influence of the GEC parameters is depicted in **Supp Figure 10**). Specifically, the bifurcation parameter for a node governs the transition between noise-dominated and oscillatory behavior. The current work assumed that strokes do not affect the bifurcation parameters/nodes, only the connections between nodes, yet prior studies indicate that delta waves are prominent in perilesional tissue and propagate to directly connected regions ^57,58^. Therefore, nodes for perilesional/partly damaged parcels and perhaps directly connected parcels may have abnormal bifurcation parameters. Evaluating this possibility is beyond the scope of this paper but is currently in progress. On the positive side, properly accounting for abnormal bifurcation parameters/nodes may improve model accuracy. On the negative side, it is unclear how node abnormality might be incorporated into a fully predictive model.

Although the GEC parameters encode directed influences between parcels, only undirected influences were assessed in the data, via Pearson correlation, and used to evaluate model accuracy. Finally, it is worth noting that stroke patients tend to be older than the usual populations where the assumptions of neurovascular coupling and the typical analysis pipelines are based ^59^. Additionally, the presented results should be interpreted given the caveat that the BOLD signal is not a direct measure of neuronal activity. Changes in network dynamics across timepoints thus reflect changes in observable BOLD fluctuations, but not necessarily nor specifically changes in neuronal dynamics.

### Conclusion

The current study shows that the effect of stroke-induced perturbations in structural connectivity on functional dynamics can be partly captured by a fully predictive whole-brain computational model, thereby demonstrating how out-of-population analyses can be used to validate whole-brain computational models. Adding lesion information to a model trained on healthy functional data is sufficient to reproduce functional anomalies in patient. Although the accuracy of prediction worsens for patients showing greater structural damage and functional deficits, the predictive model can still provide unique insights into how strokes disrupt resting brain organization.

## Acknowledgment

We thank Melina Timplalexi, Wiep Stikvoort, and Sebastian Geli for their comments and advice in the writing process.

