## Supplementary Figures for "Generative whole-brain dynamics models from healthy subjects predict functional alterations in stroke at the level of individual patients"

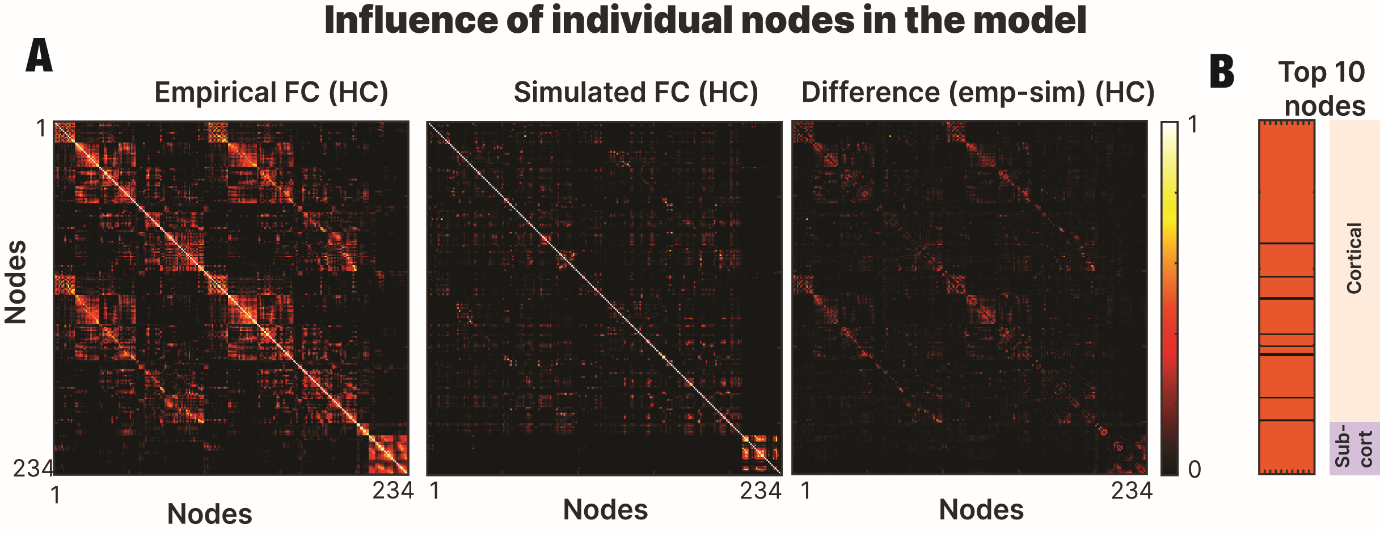
Supplementary Figure 1: Influence of connectome nodes in the model: (**A**) Using the healthy control group, we calculated the empirical FC, simulated FC and the corresponding difference between the previous two matrices. The output was a difference matrix indicating the accuracy of the fitting in each pair of nodes. (**B**) We displayed the 10 nodes with the highest difference value revealing the locations where the model was less accurate.

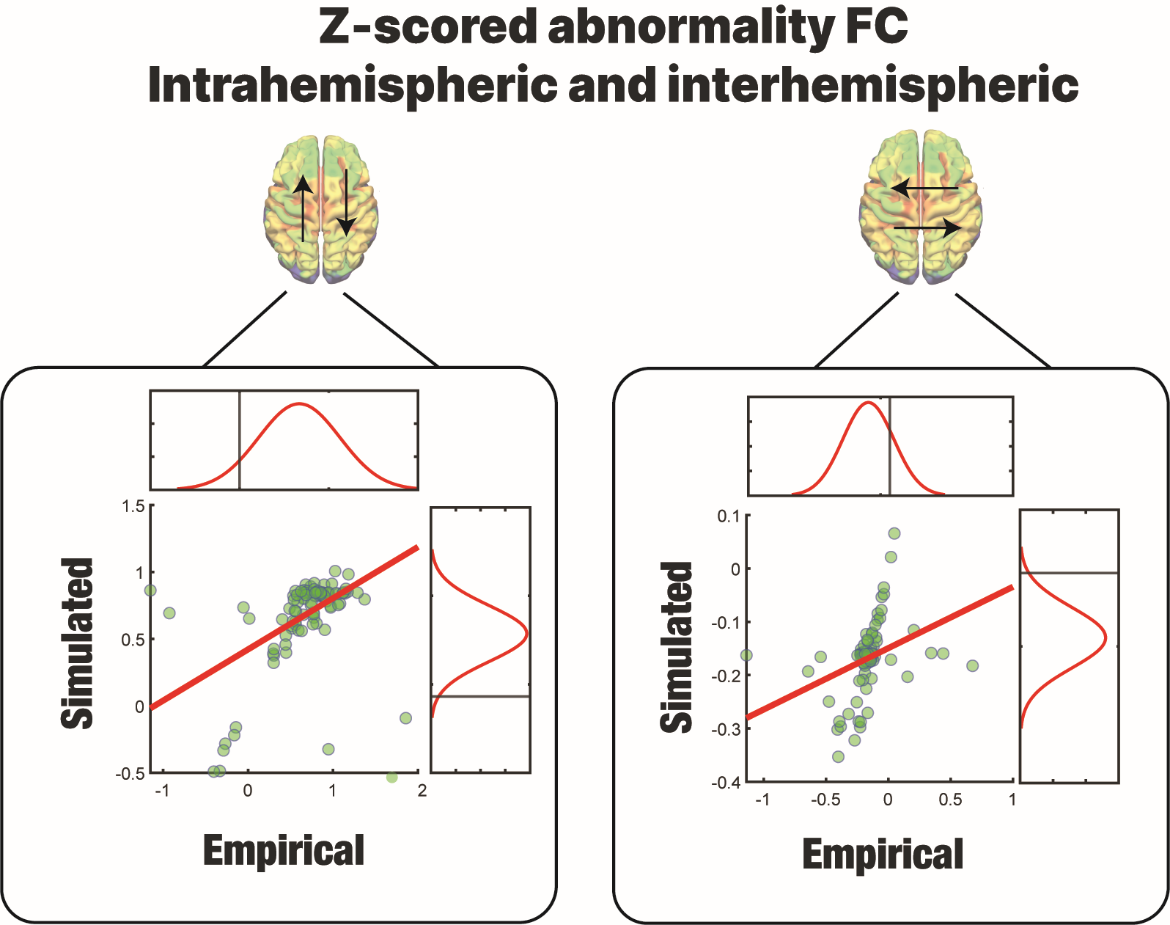

Supplementary figure 2: Model accuracy for predicting patient z-scored abnormalities in DAN-DMN FC (left panel) and homotopic interhemispheric FC (right panel). For each patient, the entries in the z-scored FC abnormality matrix corresponding to DAN-DMN FC and interhemispheric homotopic FC were separately averaged for both the empirical and simulated abnormality matrices. Then the simulated and empirical averaged quantities were plotted against one another across patients. Because DAN-DMN FC for patients tends to be more positive than controls, the corresponding patient z-scores also tends to be positive. Conversely, because interhemispheric homotopic FC for patients tends to be less positive than controls, the corresponding patient z-scores tends to be negative.

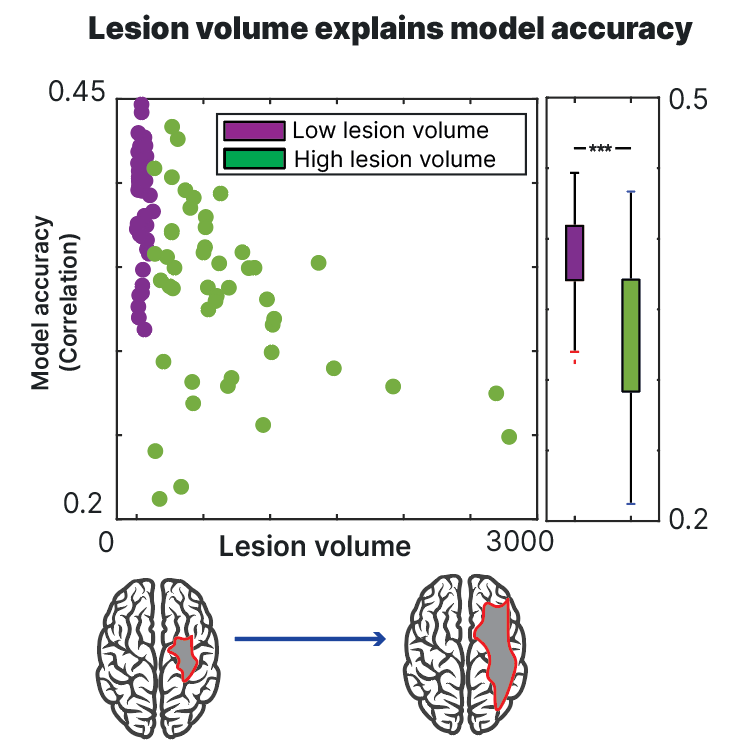

Supplementary Figure 3: Relation of lesion volume to model accuracy: Patients with larger lesion volumes showed lower model accuracies (correlation between empirical and simulated data).

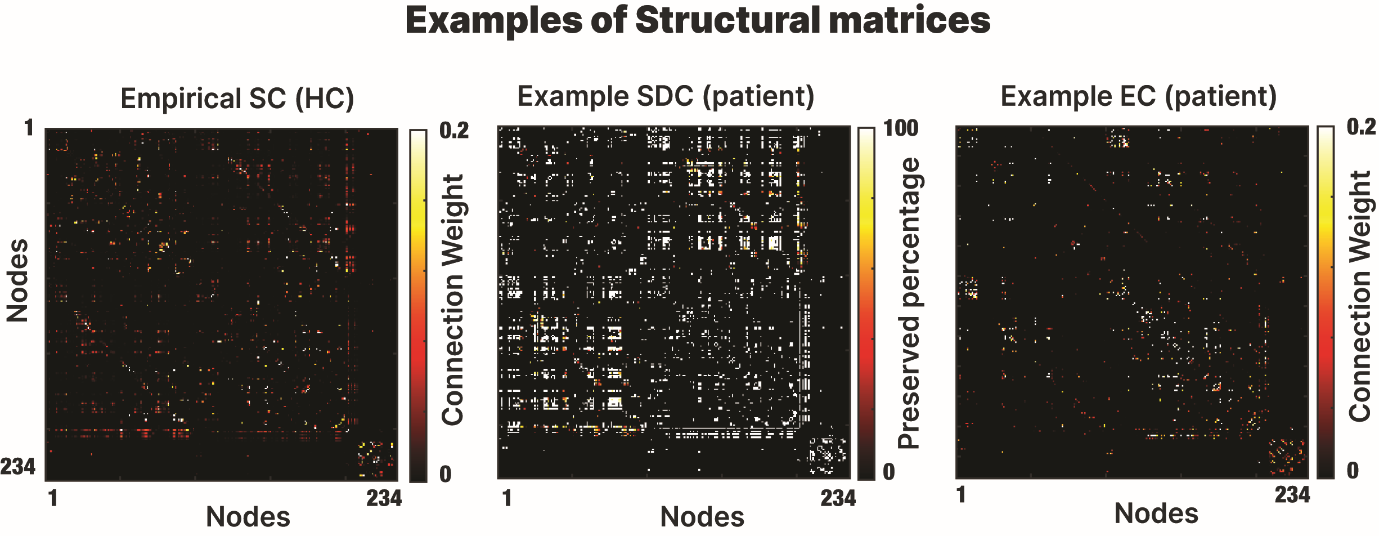

Supplementary Figure 4: Anatomical information: Matrix representation of the healthy group structural connectivity (left), SDC matrix (center) and effective connectivity matrix (right) of one stroke patient for visualization purposes.

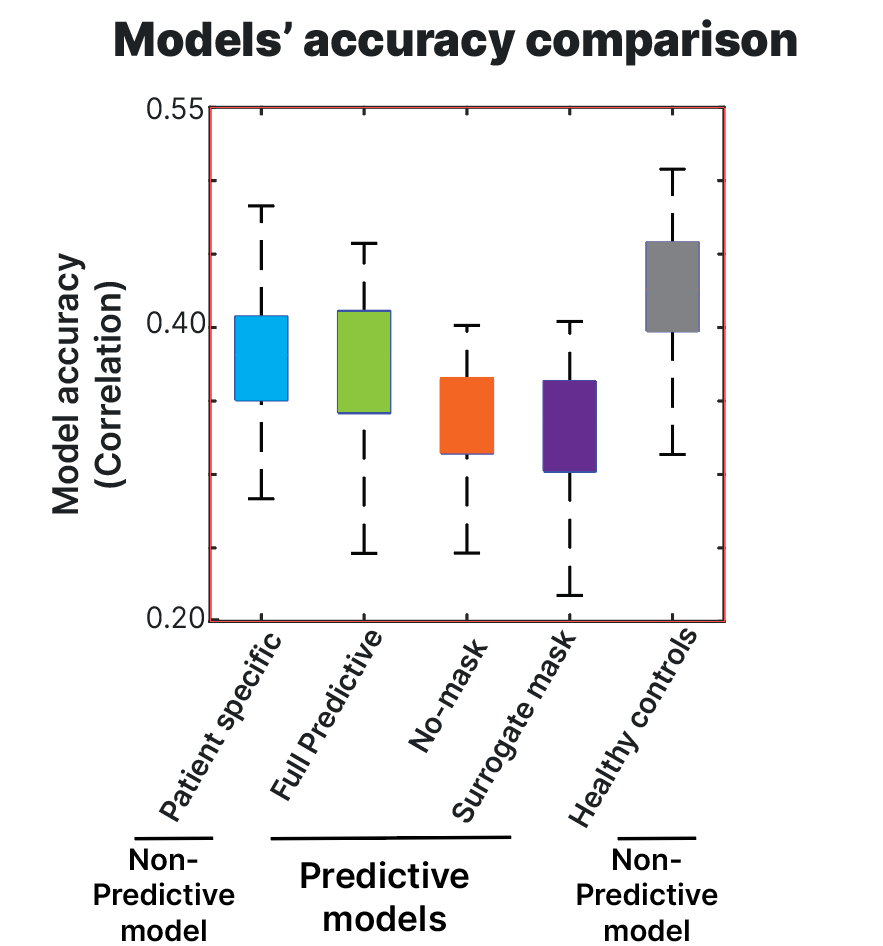

Supplementary Figure 5: Accuracy of models for patients and controls. Model accuracy was assessed by computing the correlation between the empirical and simulated FC matrices. The non-predictive patient specific model and the full predictive model showed equivalent accuracy that exceeded the accuracy for the surrogate mask model and the no-mask model. There was a significant difference between the group (*F*(4,406)= 27.84, *p*<.01) as the healthy control model showed the best accuracy and was included to assess the influence of the stroke lesion on model accuracy.

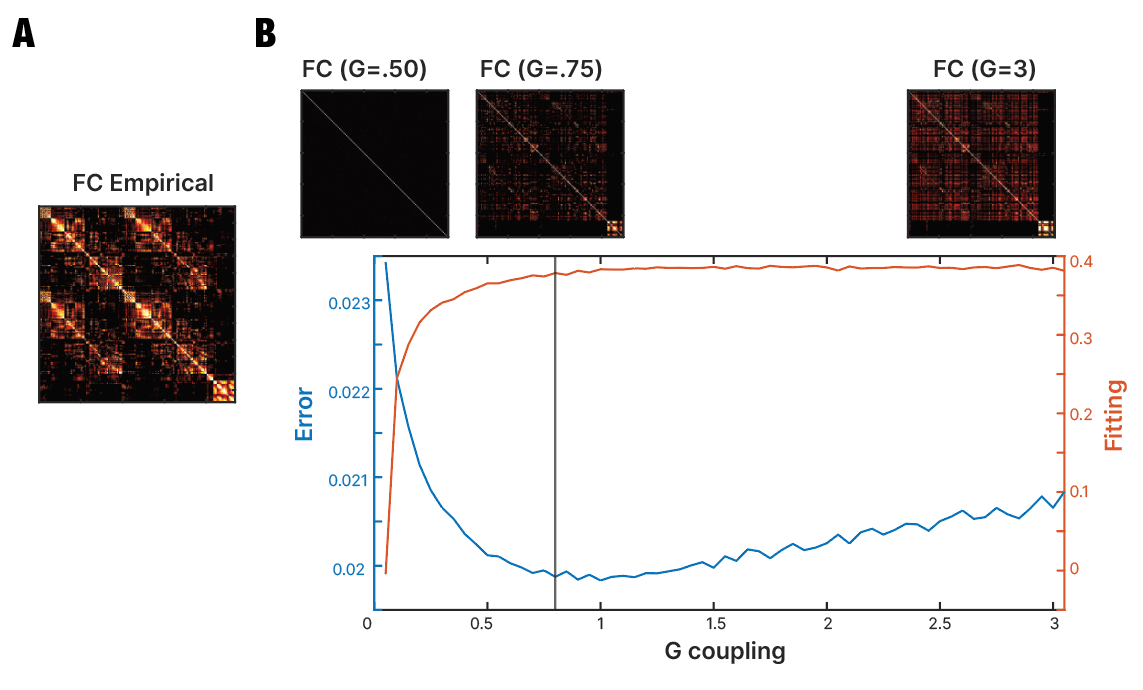

Supplementary Figure 6: Influence of GC: The similarity between the **(A)** empirical and **(B)** simulated data was assessed with different values of global coupling to determine the optimal point for the simulations.

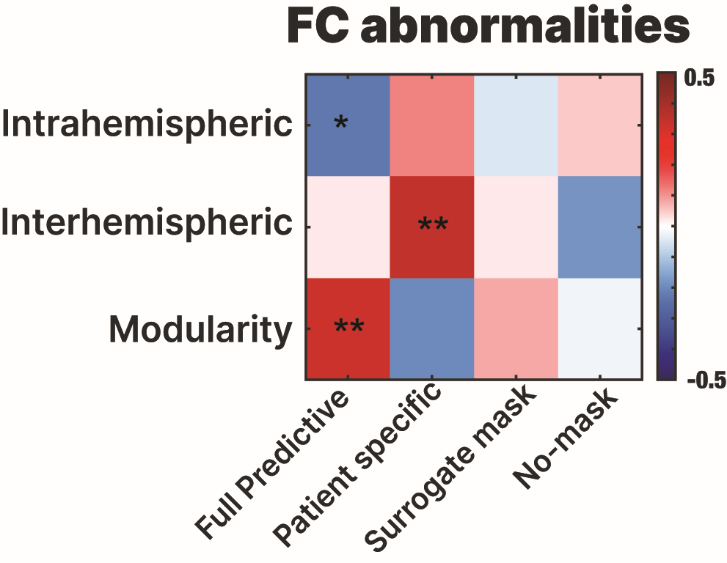

Supplementary Figure 7: Relation of model accuracy to the magnitude of FC abnormalities: The accuracy of the patient specific model was significantly correlated with inter-hemispheric FC (R=.41, p<.01), but not with intra-hemispheric FC or modularity. All the other models (including all the possibilities of the surrogate mask model and the No-mask model), showed no significant relationship with FC abnormalities.

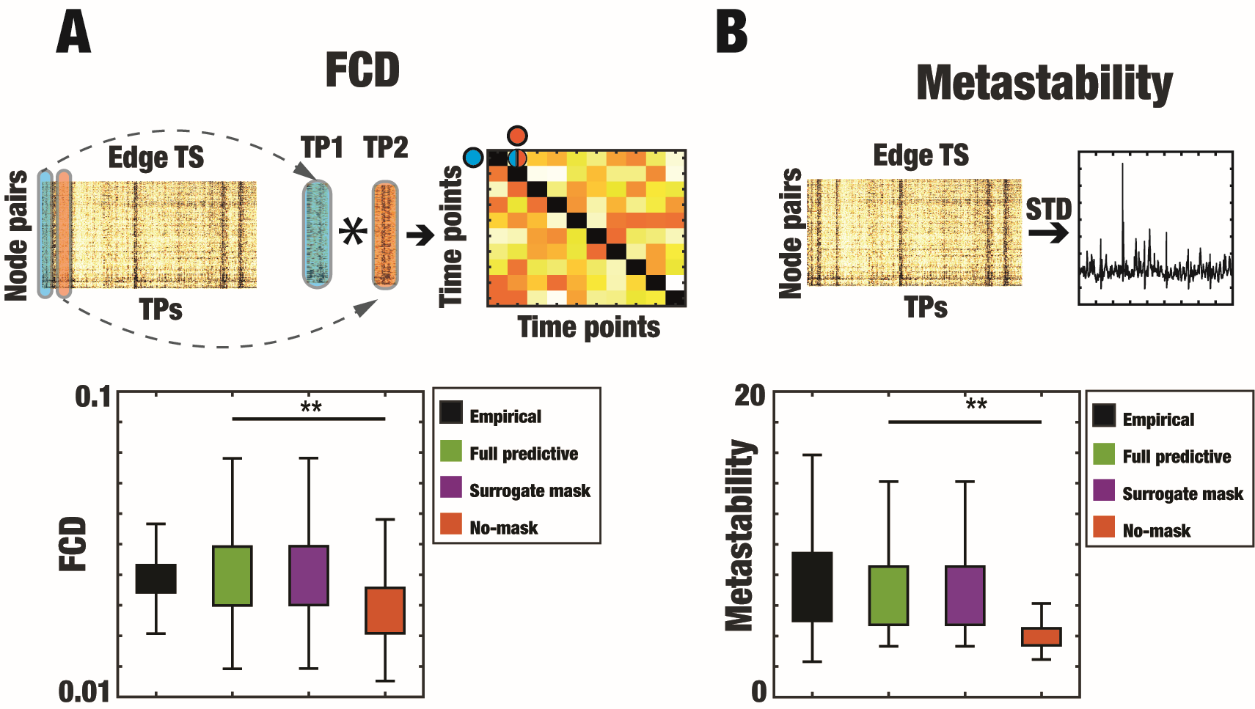

Supplementary figure 8: Dynamical metrics calculation for the different models **(A)** Time versus-time matrix representing the functional connectivity dynamics (FCD), where each entry FCD(t1, t2) is defined by a measure of resemblance between FC(t1) and FC(t2). Therefore, the FCD captures the spatiotemporal organization of FC by representing the coincidences between FC(t) matrices. It results in a symmetric matrix where an entry (ts1, ts2) is defined by the Pearson correlation between FC(ts1) and FC(ts2). FCD was estimated for the empirical data and the three predictive models showing a significant difference between the predictive models and the model without mask. Specially, there was a significant difference between the full-predictive model and the “No-mask” model (*t*(95)= 31.87, *p*<.01) **(B)** We calculated the standard deviation of the edge time series which represents the temporal metastability. This metric gives information about temporal variability in the level of synchronization. Edge metastability was estimated for the empirical data and the three predictive models showing a significant difference between the predictive models and the model without mask. In particular, the difference was significant between the full-predictive model and the “No-mask” model (*t*(95)= 19.05, *p*<.01)

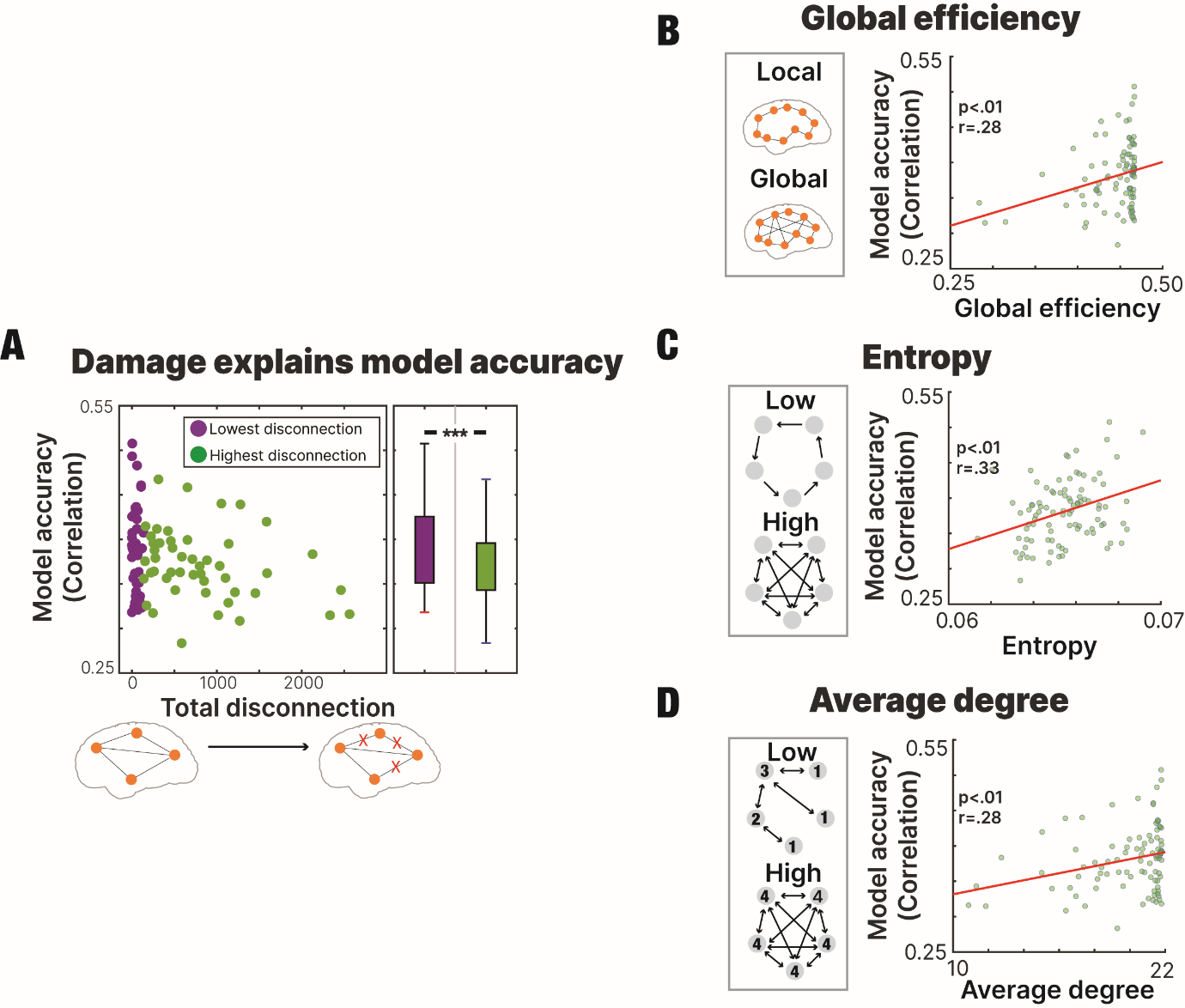

Supplementary figure 9: Relation of the accuracy of the non-predictive patient specific model to stroke metrics (**A)** Subjects with lower levels of disconnection exhibited a higher correlation between the empirical and simulated FC matrices, indicating better model performance for patients with less severe lesions. **(B-C-D)** Higher global efficiency **(B)**, higher entropy **(C)** and higher average degree **(D)** were associated with higher model accuracy.

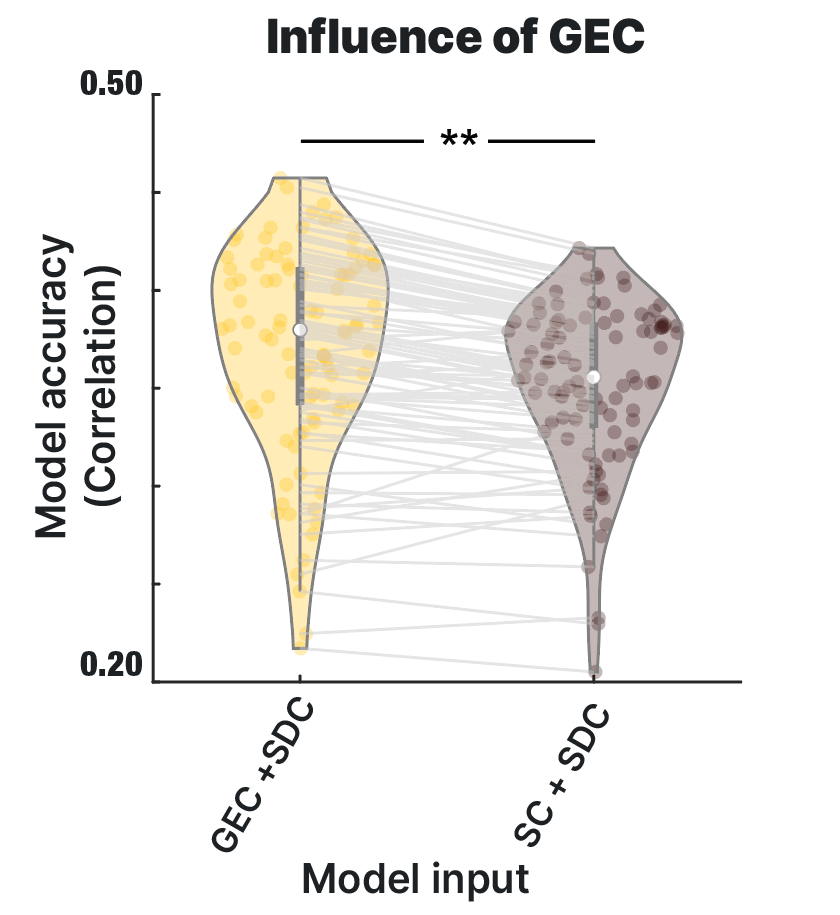

Supplementary figure 10: Influence of GEC: We compared the performance of the model when using as an input the Generative effective connectivity or the structural connectivity (both with the inclusion of the lesion disconnection mask). The GEC showed a significantly higher performance showing the relevance of model optimization.
